# Extended Bayesian inference incorporating symmetry bias

**DOI:** 10.1101/698290

**Authors:** Shuji Shinohara, Nobuhito Manome, Kouta Suzuki, Ung-il Chung, Tatsuji Takahashi, Pegio-Yukio Gunji, Yoshihiro Nakajima, Shunji Mitsuyoshi

## Abstract

In this study, we start by proposing a causal induction model that incorporates symmetry bias. This model has two parameters that control the strength of symmetry bias and includes conditional probability and conventional models of causal induction as special cases. It can reproduce causal induction of human judgment with high accuracy. We further propose a human-like Bayesian inference method to replace the conditional probability in Bayesian inference with the aforementioned causal induction model. In this method, two components coexist: the component of Bayesian inference, which updates the degree of confidence for each hypothesis, and the component of inverse Bayesian inference that modifies the model of each hypothesis. In other words, this method allows not only inference but also simultaneous learning. Our study demonstrates that the method with both Bayesian inference and inverse Bayesian inference enables us to deal flexibly with unsteady situations where the target of inference changes occasionally.

## 1 Introduction

As a cognitive bias observed in humans, the dispositions to infer “if Q, then P” and “if not P, then not Q” from “if P then Q” are well documented (Sidman and Tailby, 1982; Yamazaki, 2004; Markman and Wachtel, 1988). The former is termed symmetry bias (Takahashi et al., 2011) and the latter is termed mutual exclusivity bias (Markman and Wachtel, 1988). Consider the following example. We tend to infer “I will take you out if and only if you clean the room” and “If you don’t clean the room, then I will not take you out” from “If you clean the room, then I will take you out.” Although these inferences are invalid according to classical logic, various people are inclined to make them regardless of age.

In the field of cognitive psychology, experiments on causal induction were carried out, seeking to identify how humans evaluate the strength of causal relations between two events from their co-occurrence. In the case of a regular conditional form such as “if *p* then *q*,” the degree of confidence for the statement is considered to be proportional to the conditional probability *P*(*q* | *p*), which is the probability of occurrence of *q* following the occurrence of *P*(Evans et al., 2003). On the other hand, in case of causal relations, it has been experimentally demonstrated that humans have a strong understanding of causal relation between a cause *c* and an effect *e* when *P*(*c* | *e*) is high, as well as when *P*(*e* | *c*) is high, where *P*(*c* | *e*) is a conditional probability of the antecedent occurrence of *c*, given the occurrence of *e*. Specifically, the causal strength humans express between *c* and *e* can be approximated well by the geometric mean of *P*(*e* | *c*) and *P*(*c* | *e*). This is called the dual-factor heuristics (*DFH*) model (Hattori and Oaksford, 2007). If the causal intensity between *c* and *e* is denoted as *DFH* (*e* | *c*), then 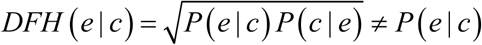. Here, note that *DFH* (*c* | *e*) = *DFH* (*e* | *c*), that is, the symmetric relationship holds.

Bayesian inference can be considered an algorithm for inferring the cause from the effect based on the notion of conditional probability. Bayesian inference speculates the hidden cause behind observations by retrospectively applying statistical inferences. The relation between Bayesian inference and brain function has been attracting attention in recent years in the field of neuroscience (Dehaene, 2014; Chater and Oaksford, 2008). In Bayesian inference, the degree of confidence in a hypothesis is updated based on a hypothetical model predefined in the form of conditional probability and current observational data. In other words, Bayesian inference is a process of narrowing down hypotheses (cause) to one that best explains observational data (effect).

Consider a situation where you estimate the emotions of others. When estimating the emotion of others, since we cannot directly observe private internal states, there is no way to estimate their emotion other than by using external clues (observational data) such as facial expression and tone of voice. In this case, emotion corresponds to a hypothesis (cause) in Bayesian inference in the sense that it is the inference target. In addition, the model corresponds to a probability distribution that represents which facial expression appears with what probability and in which emotion. For example, if you have a model such as “If she/he is pleased, there is an 80% chance that she/he will smile” and if you actually observe her/him smiling frequently, your confidence for the hypothesis that “She/he is pleased” will increase. In other words, by observing the smile (effect), you can guess the emotion of joy as being the cause of the smile.

Attention should be paid to the following two points. First, in general, in order to estimate the cause more accurately, it is better to observe more data. However, this can be said only if it is ensured that the observational data originate from the same cause. Emotions are not always constant: it is a variable that changes from time to time. For targets such as emotion, inference must be drawn as quickly as possible within a short period. Here a trade-off appears between the accuracy of estimation and the followability to changes. The second is that it is not possible to have an emotion model of the person that you meet for the first time, in advance. In such a situation, it is necessary that the model is learned simultaneously with the inference being drawn. If this model is wrong, the correct reasoning for the emotion of the person cannot be acquired.

Arecchi (2011) proposed the concept of the inverse Bayesian inference where the model is modified according to circumstances. Gunji et al. (2018) and Horry et al. (2018) formulated the inverse Bayesian inference and demonstrated that animal herding and human decision-making can be satisfactorily modeled by combining Bayesian inference and inverse Bayesian inference. This framework can be said to simultaneously perform both Bayesian inference that picks up the optimal hypothesis from the predefined set of hypotheses, and inverse Bayesian inference that creates or modifies the models of hypotheses based on observational data. The latter can be said to be a learning of the model, rather than an inference.

In this paper, first, we propose a causal induction model that incorporates symmetry bias by parametrizing the mixing rate of *P*(*e* | *c*) and *P*(*c* | *e*), i.e., the strength of symmetry bias. Second, we propose a human-like Bayesian inference where the conditional probability schema in Bayesian inference is replaced with the causal induction model proposed above. We term the inference “extended Bayesian inference” in the sense that it includes Bayesian inference as a special case where the strength of symmetry bias is zero. Finally, we conducted the simulation, derived from the problem of inference the probability of getting heads in the course of repetitive coin toss and verified the validity of the extended Bayesian inference.

## 2 Material and methods

### 2.1 Causal induction models

This section describes simple causal induction models that infer the strength of a causal relation between *c* and *e* from four pieces of co-occurrence concerning the cause candidate *c* and the effect event *e* (Table 1). The most representative model of causal induction is the Δ*P* model (Jenkins and Ward, 1965). It takes the difference between the conditional probability of occurrence of *e* given the occurrence of *c* and the conditional probability of occurrence of *e* given non-occurrence of *c* as an index for causal strength.

**Table 1.** The 2 × 2 contingency table for elemental causal induction.

| | effect ($e$) | no effect ($\neg e$) | marginal frequency |
| --- | --- | --- | --- |
| cause ($c$) | $N(c, e)$ | $N(c, \neg e)$ | $N(c)$ |
| no cause ($\neg c$) | $N(\neg c, e)$ | $N(\neg c, \neg e)$ | $N(\neg c)$ |
| marginal frequency | $N(e)$ | $N(\neg e)$ | |

$N(x)$ denotes the frequency of occurrence of $x$ and $N(x, y)$ denotes the frequency of co-occurrence of $x$ and $y$ .

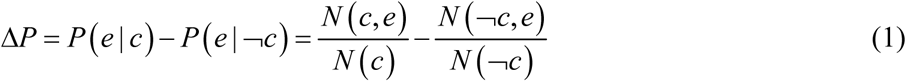

This model formalizes the basic idea of scientific experiments. For example, the first term, *P*(*e* | *c*), represents the test group whereas the second term, *P*(*e* | ¬*c*), represents the control group (Takahashi et al., 2007).

Hattori and Oaksford (2007) proposed the *DFH* model, which showed the best fit compared to 32 models without parameters and eight models with parameters. This model is based on the geometric mean of, which stands for the predictability of the effect from the cause, and its inverse *P*(*c* | *e*).

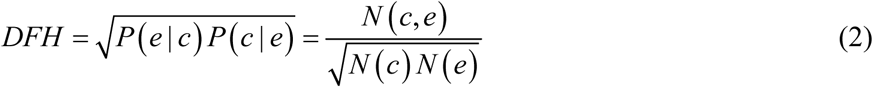

The strength of relevance between rows and columns in the 2 × 2 table such as Table 1 is expressed using the four-fold correlation coefficient *ϕ*.

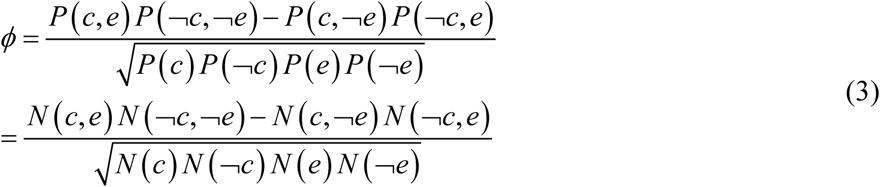

This formula includes *N* (¬*c*, ¬*e*), which represents the frequency of the co-occurrence of the absence of *c* and the absence of *e*. In general, the event wherein something does not occur is difficult to define clearly and enumerate, and the frequency of the co-occurrence of such the events may be innumerable. *DFH* is expressed as the limit where *N* (¬*c*, ¬*e*) in *ϕ* is infinite (Hattori and Oaksford, 2007).

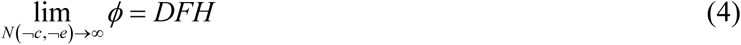

In other words, *DFH* can be said to be a simplified version of *ϕ* that can be used under the situations where the occurrences of *c* and *e* are extremely rare compared to the absences of *c* and. *e*.. As for Δ*P*, if *N* (¬*c*, ¬*e*) → ∞ then *N* (¬*c*) → ∞, so the following holds:

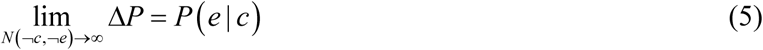

In other words, Δ*P* approximates the conditional probability (Takahashi et al., 2007).

Aside from the Δ*P* model and *DFH* model, Takahashi et al. (2007) proposed the proportion of assumed-to-be rare instances (*pARIs*) as yet another model that has an unusually high affinity with the human causal induction judgment.

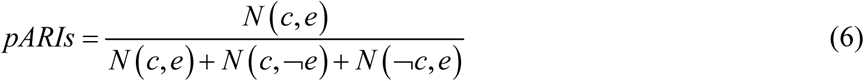

As described above, the *DFH* model is defined as the geometric mean of *P*(*e* | *c*) and *P*(*c* | *e*). *pARIs* can be transformed as follows.

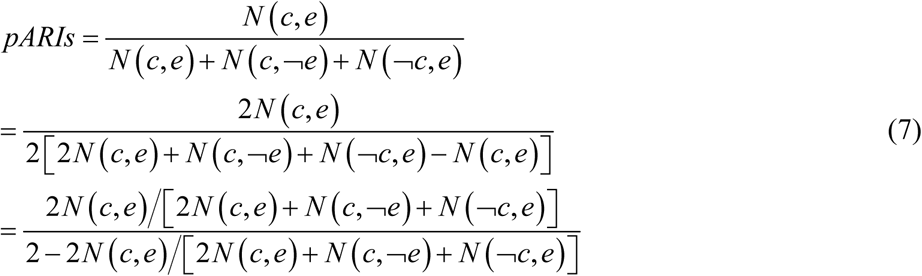

On the other hand, the harmonic mean (*HM*) of *P*(*e* | *c*) and *P*(*c* | *e*) can be described as follows.

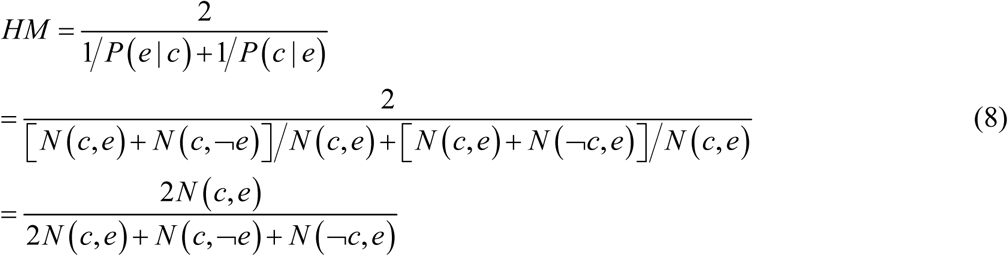

Therefore, *pARIs* and *HM* are related by monotonically increasing functions that are in one-to-one correspondence.

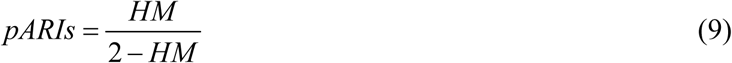

According to this, *pARIs* can be seen as a disguised form of *HM*. Thus, *DFH* and *pARIs* have similar biconditional forms (“if *c* then *e*, and if *e* then *c*”), although there is a difference between geometric means and *HM* (Takahashi et al., 2007).

If *pARIs* is expressed as *pARIs* (*e* | *c*) and *c* in the formula (6) is replaced by *e*, we get:

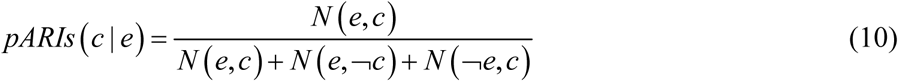

Because *N* (*x, y*) = *N* (*y, x*), we ultimately get

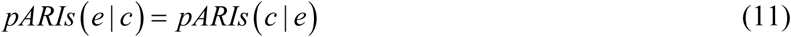

and the symmetry obtains as in *DFH*.

### 2.2 Extended confidence model

Consider a generalized weighted average of *P*(*e* | *c*) and *P*(*c* | *e*) as a biconditional causal induction model that includes both *P*(*e* | *c*) and *P*(*c* | *e*). First, the generalised weighted average of two variables, *x* and *y* can be expressed in the following formula using parameters *α* and *m*

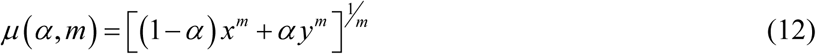

Where *α* takes values in the range 0.0 ≤ *α* ≤ 1.0 and denotes a weighted value of *x* and *y*, and *m* takes values in the range −∞ ≤ *m* ≤ ∞ and denotes the kind of the mean. For example, suppose *α* = 0.5 and *m* = 1.0, then *μ* (0.5,1.0) = 0.5*x* + 0.5 *y* representing the arithmetic mean. Suppose *α* = 0.5 and *m* = −1.0, then *μ* (0.5, −1.0) = 2*xy* (*x* + *y*) representing the harmonic mean. Supposing *m* = 0.0, the formula (12) is undefinable. If, however, we represent the mean value in the limit of *m* → 0.0 as *μ* (*α*, 0.0), we get *μ* (*α*, 0.0) = *x*^1−*α*^ *y*^*α*^, where if *α* = 0.5, the geometric mean 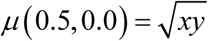.

Here, we express the connection strength between two events *c* and *e* as *C*(*e* | *c*). Hereinafter, this will be termed the Extended Confidence Model.

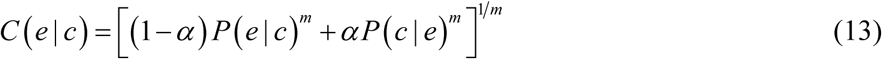

Here, the parameter *α* can be regarded as a parameter that controls the strength of the symmetry bias which takes maximum value when *α* = 0.5. When *α* = 0.0, *C*(*e* | *c*) = *P*(*e* | *c*) obtains irrespectively of the value of *m*, and *C*(*e* | *c*) expresses a normal conditional probability. If *α* = 0.5 and *m* = 0.0, 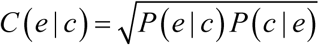 obtains and it coincides with the *DFH* model. In addition, *pARIs* is associated with *C*(*e* | *c*) in the case of *α* = 0.5 and *m* = −1.0. Thus, the proposed model can be described as an extended model which accommodates the conditional probability *P*(|), *DFH* and *pARIs* as special cases.

As shown in the appendix, we derived the values of *α* and *m* that best fit the causal strength judged by humans using equation (13) and the eight types of experimental data shown in the literature (Hattori and Oaksford, 2007; Anderson and Sheu, 1995; Buehner et al., 2003; Lober and Shanks, 2000; White, 2003). Thus, the value of *α* was 0.35. In other words, it means that human do not evaluate *P*(*e* | *c*) and *P*(*c* | *e*) equally, but rather attach a bit more importance to *P*(*e* | *c*). On the other hand, the value of *m* was −1.15, which is close to the *HM* (*m* = −1).

### 2.3 Bayesian Inference

This study deals with the problem of inferring a generative model (probability distribution) from observational data. To this end, in what follows, the hypothesis *h* and data *d* will be used on behalf of *c* and *e*. Moreover, discrete models are considered. Bayesian inference first defines several hypotheses {*h*_*i*_} and provides a model for each hypothesis (probability distribution of data) in the form of conditional probability *P*(*d* | *h*_*i*_). When data are fixed and *P*(*d* | *h*_*i*_) regarded as a function of hypotheses, it is termed likelihood. The confidence *P*(*h*_*i*_) for each hypothesis is given as a prior probability. We can take *P*(*d* | *h*_*i*_) and *P*(*h*_*i*_) as initial values and calculate the posterior probability *P*(*h*_*i*_ | ***d***) when observing data ***d*** using Bayes’ theorem as follows.

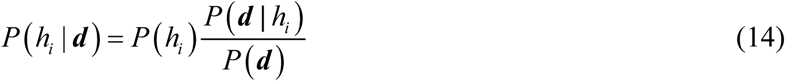

Hereinafter, data observed at a point in time are represented by the bold ***d***. Afterwards, we can replace the posterior probability with the prior probability using Bayesian updating.

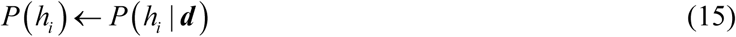

By combining formulas (14) and (15), we get

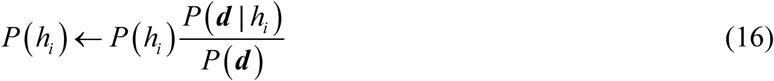

Whenever new data ***d*** are observed, *P*(*h*_*i*_) in the formula (16), i.e., confidence in each hypothesis, is updated.

The inference distribution during this procedure can be expressed as

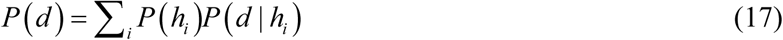

Figure 1 (a) shows an overview of the processing flow of Bayesian inference.

**Fig. 1.**
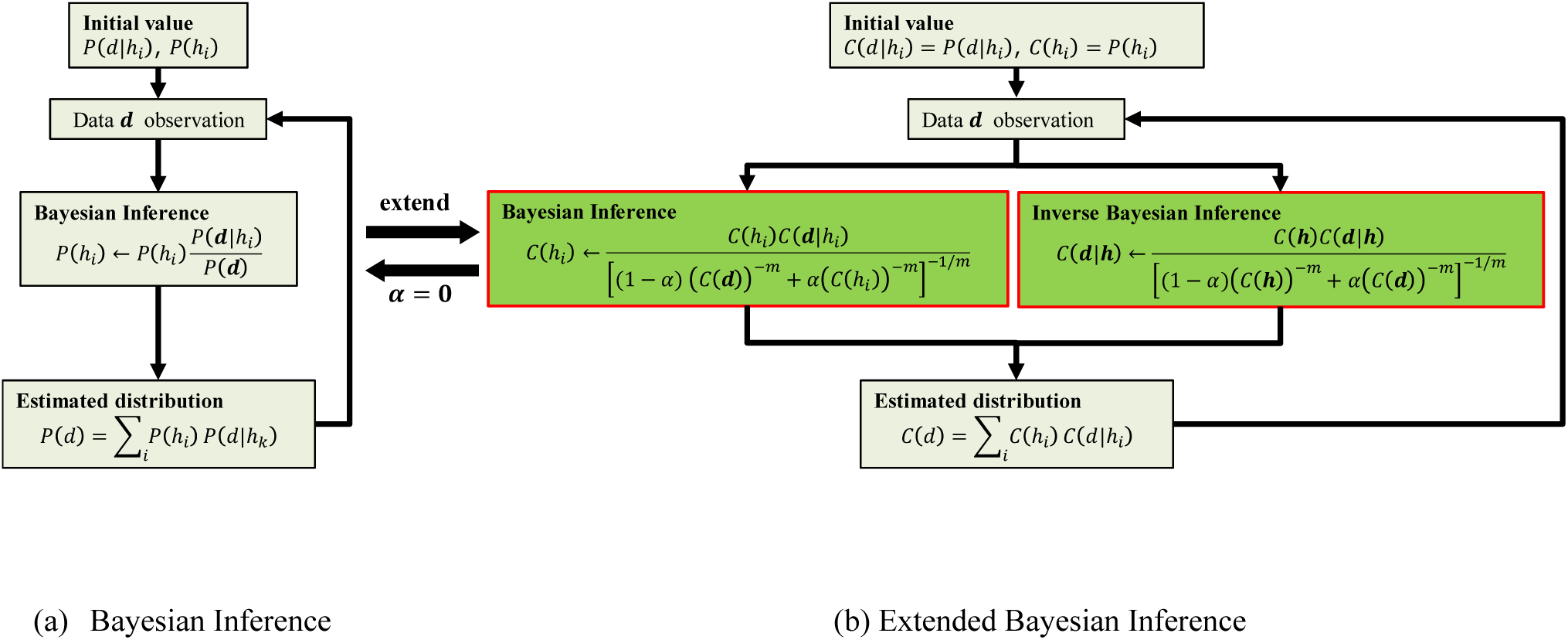
A flow chart comparing extended Bayesian inference to Bayesian Inference. (a) An overview of the processing flow of Bayesian inference; and (b) An overview of the processing flow of extended Bayesian inference. In (b), for simplicity, the portion that belongs to the normalisation process is omitted. In (b), if we suppose *α* = 0, the portion of inverse Bayesian inference disappears, corresponding to the Bayesian inference in (a).

### 2.4 Extended Bayesian Inference

In this section, we propose the extended Bayesian inference where the conditional probability in Bayesian inference is replaced with the extended confidence model. The extended Bayesian inference is an inference that has *α* and *m* as parameters and embraces normal Bayesian inference as its special case when *α* = 0. First, we can replace *c* and *e* in formula (13) with *h*_*i*_ and *d*.

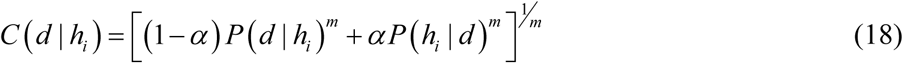

Then we apply the Bayes’ theorem to the right side of the formula (18), and we can obtain the conversion as follows.

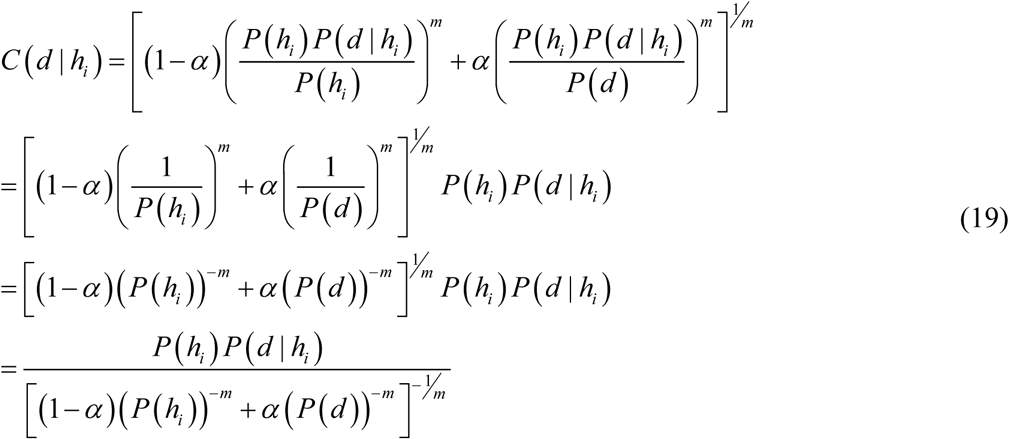

By replacing *h*_*i*_ and *d* in formula (19), to perform the same conversion, we get

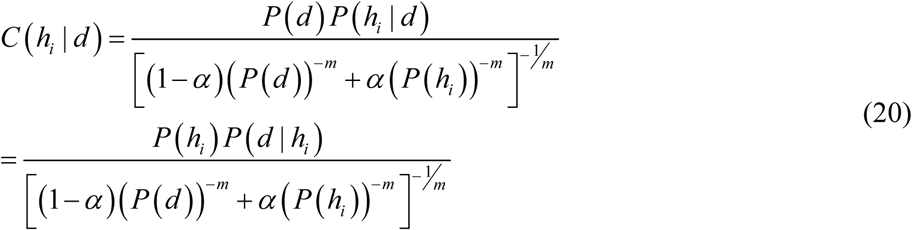

In the next step, we replace the conditional probability *P* on the right side of formulas (19) and (20) with the extended confidence *C* to make the formulas recursive, and then we replace the equation with the update formula.

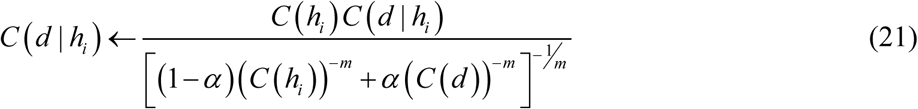

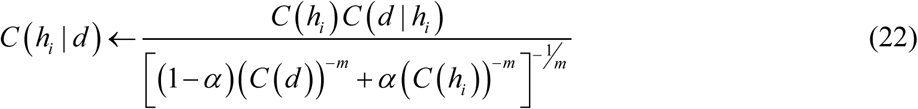

In what follows we show the processing flow of extended Bayesian inference, summarized in Fig.1(b). First, we take *P*(*d* | *h*_*i*_) and *P*(*h*_*i*_) as initial values and substitute them with *C*.

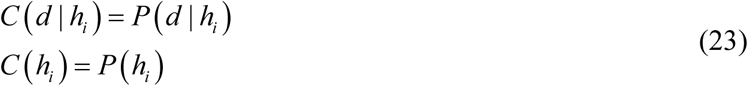

Second, we calculate the posterior probability *C*(*h*_*i*_ | ***d***) using the formula (22) when ***d*** is observed. Finally, we replace the posterior probability with the prior probability in just the same way done in Bayesian inference.

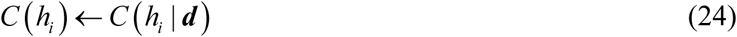

Taken together, we get

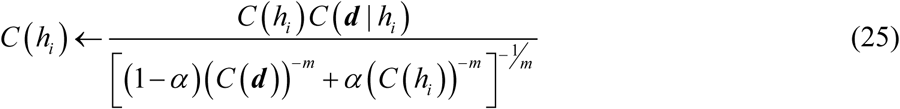

Here, we can see that, supposing *α* = 0, the right side shows the same form as that of Bayesian inference seen in formula (16).

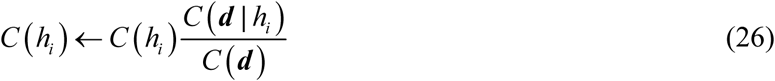

Now, in the case of Bayesian inference, the model *P*(*d* | *h*_*i*_) was invariable. We can ask if this is the same for extended Bayesian inference. Here, if we look closely to the right side of formula (21), supposing *α* = 0, formula (21) becomes a tautology as shown below.

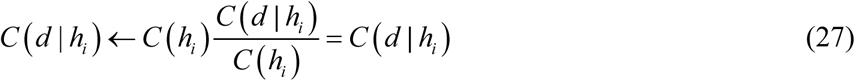

In other words, if *α* = 0, then formula (21) substantially disappears. Conversely, if *α* > 0, *C*(*d* | *h*_*i*_), is subject to the denominator *C*(*d*) in the right side, that is, the estimated value of the data. Following Gunji et al. (2016) and Gunji et al. (2018), the process shown in formula (21) is termed Inverse Bayesian Inference. As seen in formula (21), in inverse Bayesian inference, the amount of modification to the model of each hypothesis increases as *α* becomes larger. In this paper, we do not update the likelihood of all hypotheses, but update only the likelihood of the hypothesis with the highest confidence at that time. That is, the following formula is used instead of formula (21).

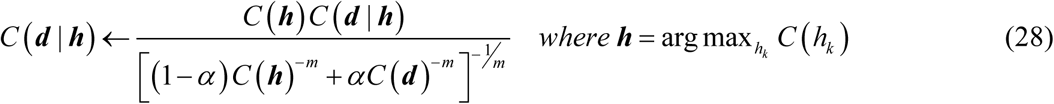

Hereafter, the hypothesis with the highest confidence is denoted as the bold ***h***. If there are multiple hypotheses with the highest confidence, one of them is selected at random.

Following the application of formulas (25) and (28), we can normalize the degree of confidence for each hypothesis and the model.

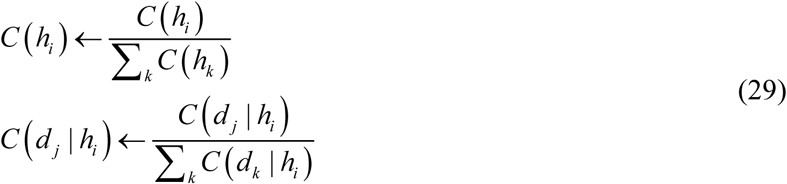

Finally, we can calculate the estimated distribution values as with Bayesian inference.

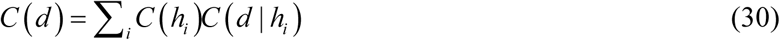

Figure 1(b) shows an overview of the processing flow of extended Bayesian inference.

### 2.5 Simulations

To observe the difference in behaviour between Bayesian inference and extended Bayesian inference, a simulation was performed. Specifically, a coin was tossed repeatedly to observe the results and estimate the probability of getting heads using both Bayesian inference and extended Bayesian inference. In the simulation, the probability of it landing heads at the *t*^*th*^ trial was designated *p*^*t*^ and the probability of it landing tails was designated as 1− *p*^*t*^ to handle cases where the probability changes over time. In each trial, a uniformly distributed random number is generated from interval [0.0, 1.0]. If the number is predefined *p*^*t*^ or more, it is regarded as getting head, if it is less than *p*^*t*^, it is regarded as getting tail.

First, let heads be expressed as *H* and tails as *T*, i.e., *d* = {*H, T*}. Second, we prepare *N* + 1 hypotheses {*h*_0_, *h*_1_, …, *h*_*N*_} and define the probability of heads and the probability of tails in each hypothesis *h*_*i*_ as follows.

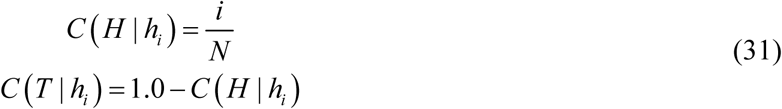

Further, we must suppose that the prior probability for each hypothesis is equal.

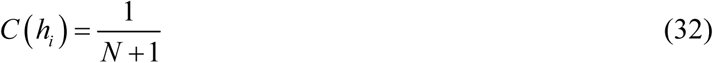

We then take these as initial values and use formulas (25), (28), (29) and (30) whenever a coin toss result is observed, in order to perform extended Bayesian inference and update the estimated value of the probability of head landing sequentially, i.e., *C*(*H*). Note that Bayesian inference is calculated on the premise that *α* = 0.0.

This article deals with the three tasks below. The first task is a case where the probability of head landing is fixed (call it Task I).

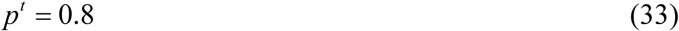

The second task is a case where the probability of head landing changes continually from 0 to 1 according to sin function (call it Task II).

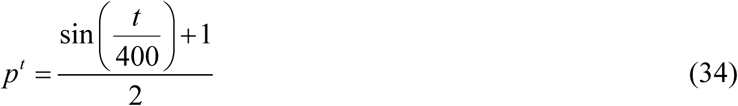

In the simulations for both Tasks I and II, *N* = 10, that is, the number of hypotheses was 11. The third task is alike the second one, differing from it in using only those hypotheses where the probability of head landing is from 0 to 0.5, i.e., cases where only six hypotheses {*h*_0_, *h*_1_,…, *h*_5_} were used for inference (call it Task III). For this reason, coins with the probability of head landing > 0.5 are unknown objects that do not exist in the system.

In formula (16), if *P*(*h*_*i*_) becomes zero at once, it always becomes zero thereafter. In order to prevent this, normalization processing (smoothing) is performed by adding a small positive constant *ε* to the confidence of each hypothesis obtained by formula (16).

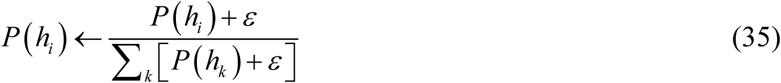

## 3 Results

This section describes the simulation results of tasks I, II and III. For the simplicity of subsequent analysis, in the following simulations, the parameter *m* was fixed to −1 in the extended Bayesian inference. Figure 2 (a), (c) shows the results of Bayesian inference and extended Bayesian inference in Task I.

**Fig. 2.**
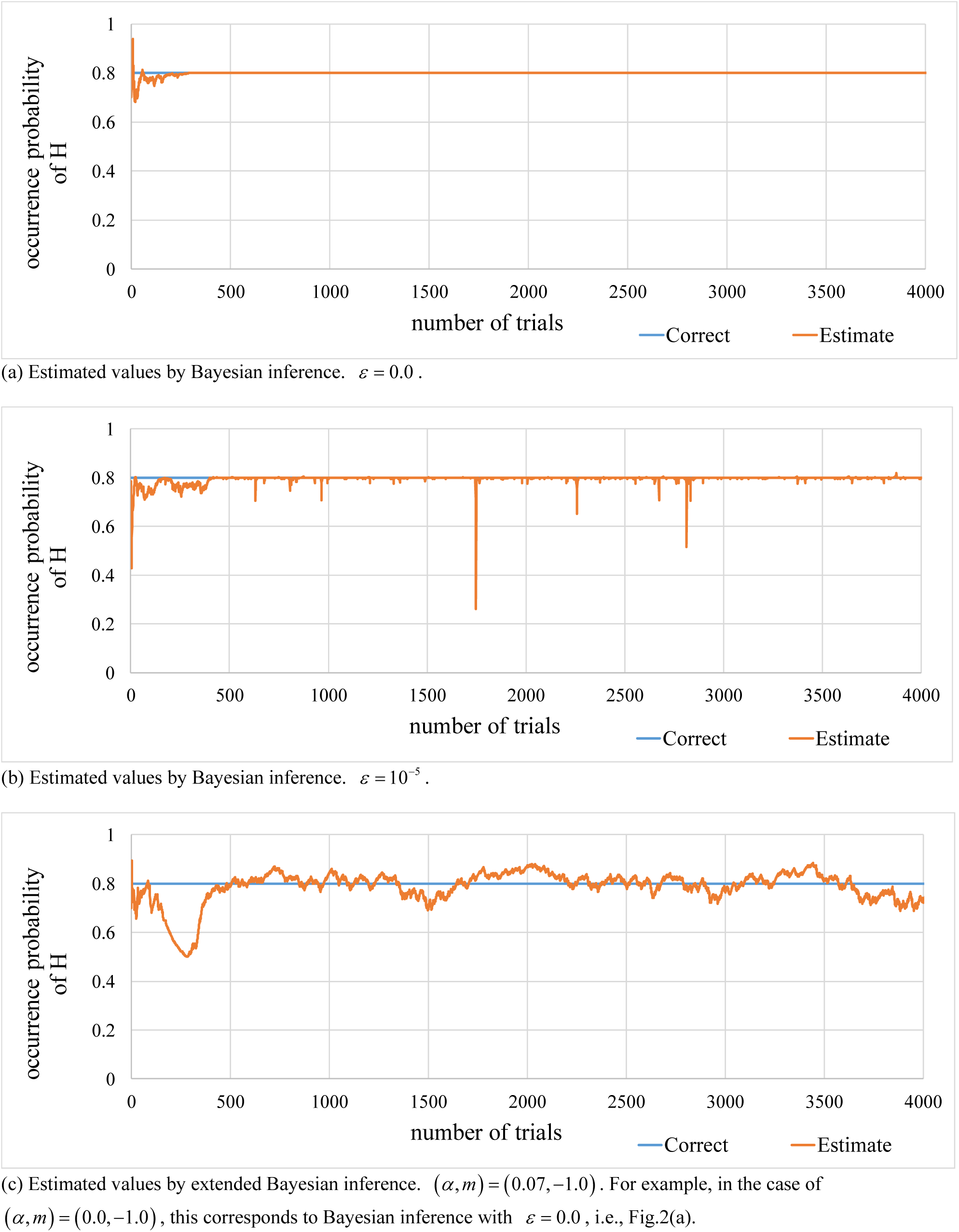
Time progress of the estimated values for the probability of head in Task I. The figure includes the correct probability.

In Bayesian inference, the estimated value quickly converges to 0.8, which is the correct probability as shown in formula (33). On the other hand, in extended Bayesian inference, the estimated value fluctuates such value. Figure 3 (a), (c) shows the results of Bayesian inference and extended Bayesian inference in Task II. Although Bayesian inference initially seemed inclined towards following the sin wave, the correct answer, it gradually declined and converged to 0.5. Regarding formula (16), representing Bayesian updating, when *P*(***d***) is viewed as a common constant across all *h*_*i*_, it can be written as follows.

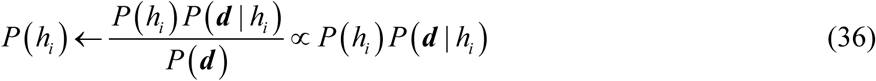

**Fig. 3.**
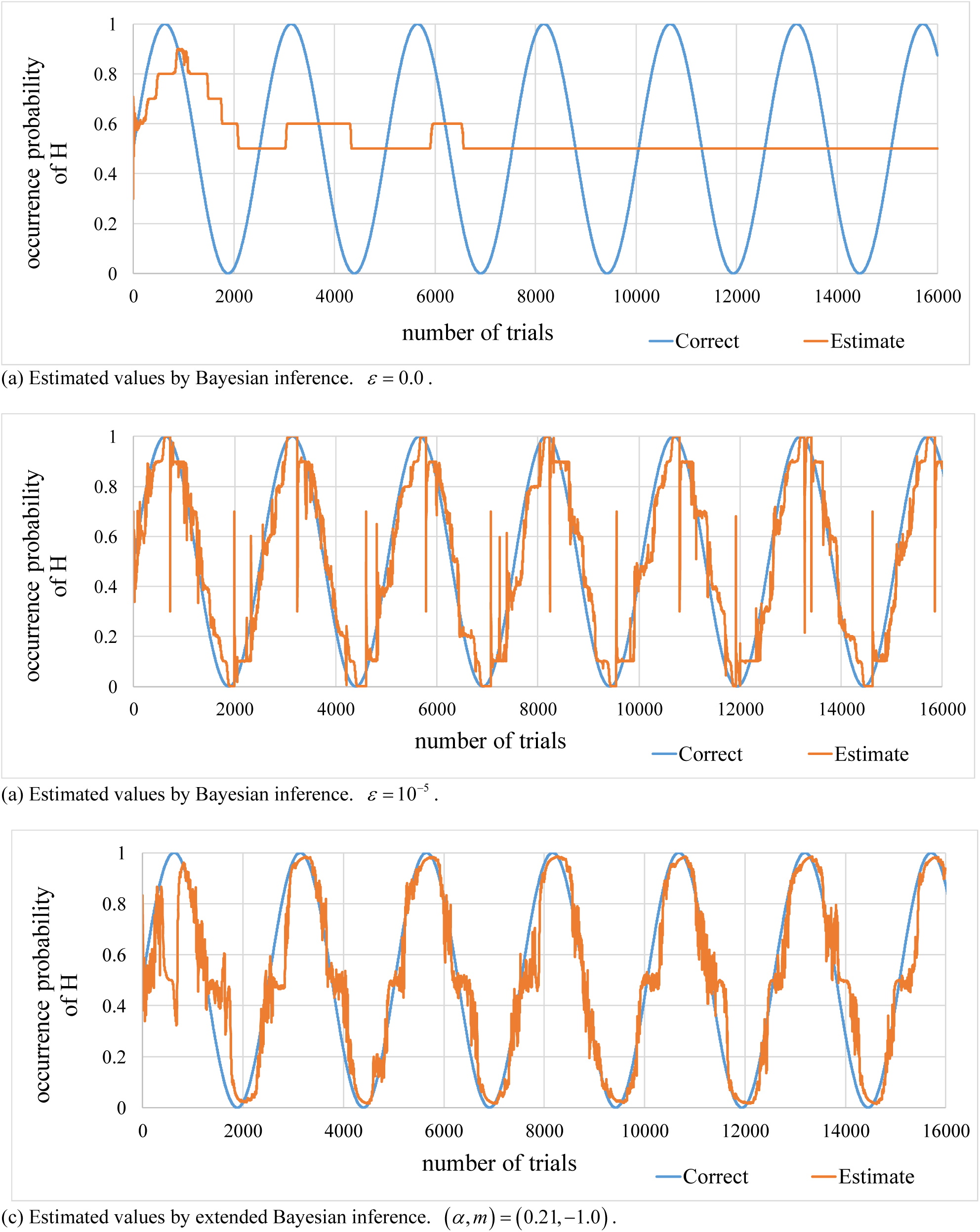
Time progress of the estimated values for the probability of head in Task II. The figure includes the correct probability.

Thus, in the case of Bayesian inference, the present degree of confidence multiplied by the likelihood for the current observational data can be used to calculate the next degree of confidence. By repeating this procedure recursively, the degree of confidence comes to reflect the observation history. For this reason, the estimated value for the probability of head landing in the Bayesian inference in Task II converges to 0.5, the average time of the correct probability (sin waves) of head landing, as expressed in formula (34).

It should be noted, however, that Bayesian inference can still handle unsteady circumstances by ignoring the observation history of the remote past, that is, in the case of *ε* > 0.0. The results of Bayesian inference when *ε* = 10^−5^ for Tasks I and II are shown in Figure 2(b) and Figure 3(b) respectively. When *ε* = 10^−5^, the Bayesian inference does not have the same functions of the extended Bayesian inference. The hypothesis model is invariable and can follow sin wave by changing the hypothesis itself.

Figure 4 shows the time progress of the estimated value for the probability of head landing in Task III. In the case of extended Bayesian inference, although the hypotheses were restricted to those with a probability of head landing 0.5 or lower, on default, we can see that it successfully follows when the probability of head landing is greater than 0.5. On the other hand, in the case of Bayesian inference, we can say that it failed to follow successfully since it did not have the same hypothesis when the correct probability of head landing was greater than 0.5.

**Fig. 4.**
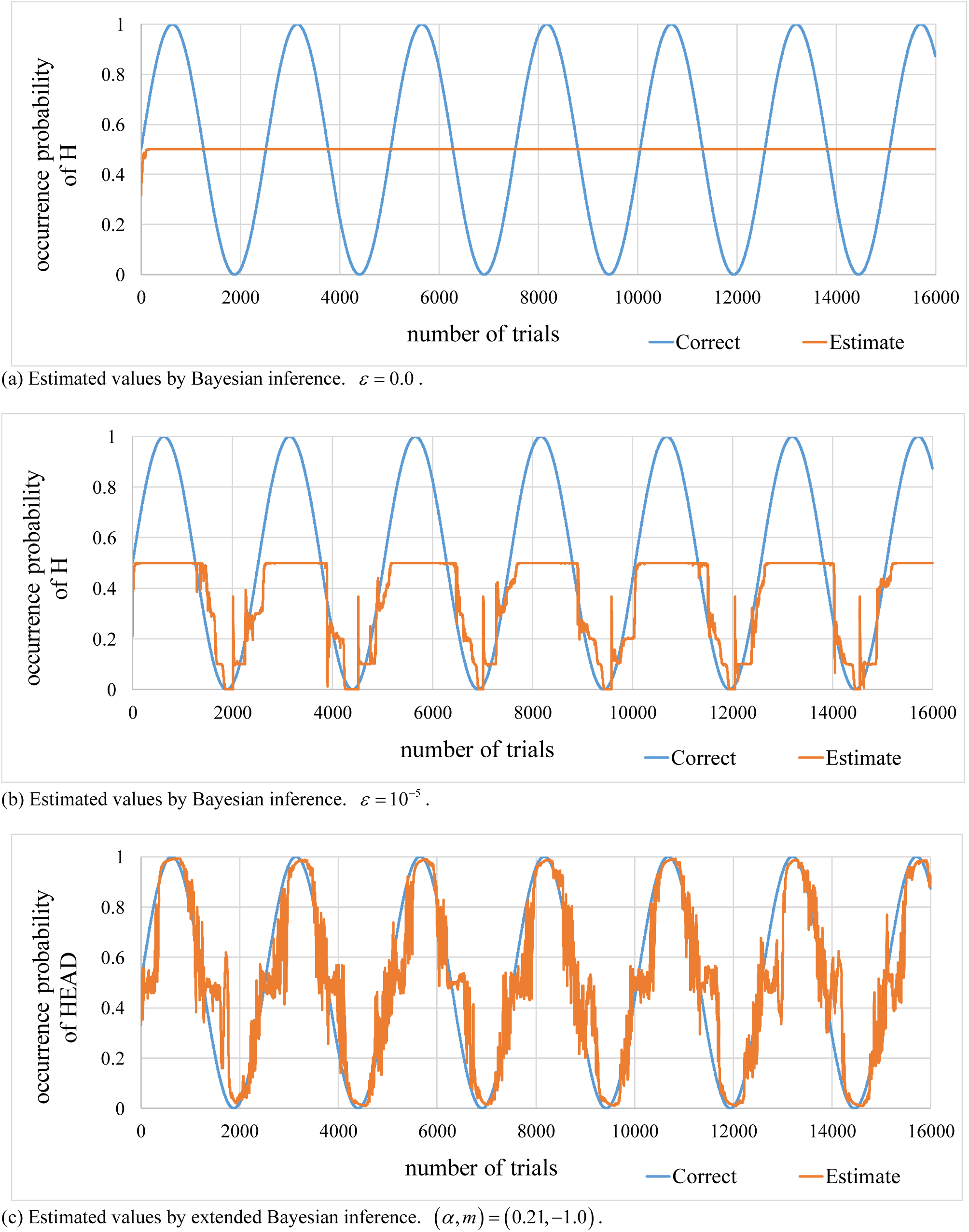
Time progress of the estimated values for the probability of head in Task III. The figure includes the correct probability.

Figure 5 shows the internal state of the extended Bayesian inference in Task II. Figure 5(a)(b) shows the time progress of the degree of confidence for each hypothesis and the hypothesis number that shows the greatest degree of confidence. These figures illustrate a repeated cycle of a stable period where the degree of confidence in a certain hypothesis reaches approximately 1 and the degree of confidence in other hypotheses reaches approximately 0. An instable period is also shown where no such hypothesis reaching 1 exists and the hypothesis with the greatest degree of confidence rapidly changes. In addition, the hypothesis with the highest degree of confidence does not always remain the same leaving the spot open, alternating amongst various hypotheses. Figure 5(c) shows the time progress of the probability of head landing for each hypothesis. It illustrates how, during the stable period, the degree of confidence in the hypothesis with the greatest degree of confidence remains unchanged at approximately 1, whilst the model follows the sin wave by changing its content. Figure 6, in general, refers to the internal state of the Bayesian inference in Task II. Figure 6(a)(b) illustrates the time progress of the degree of confidence for each hypothesis and the hypothesis number that has the greatest degree of confidence. Part (c) of Figure 6 shows the time progress of the probability of head landing for each hypothesis. With these figures, we can see how for the Bayesian inference, the models are invariant and follow the sin wave by changing the hypothesis itself.

**Fig. 5.**
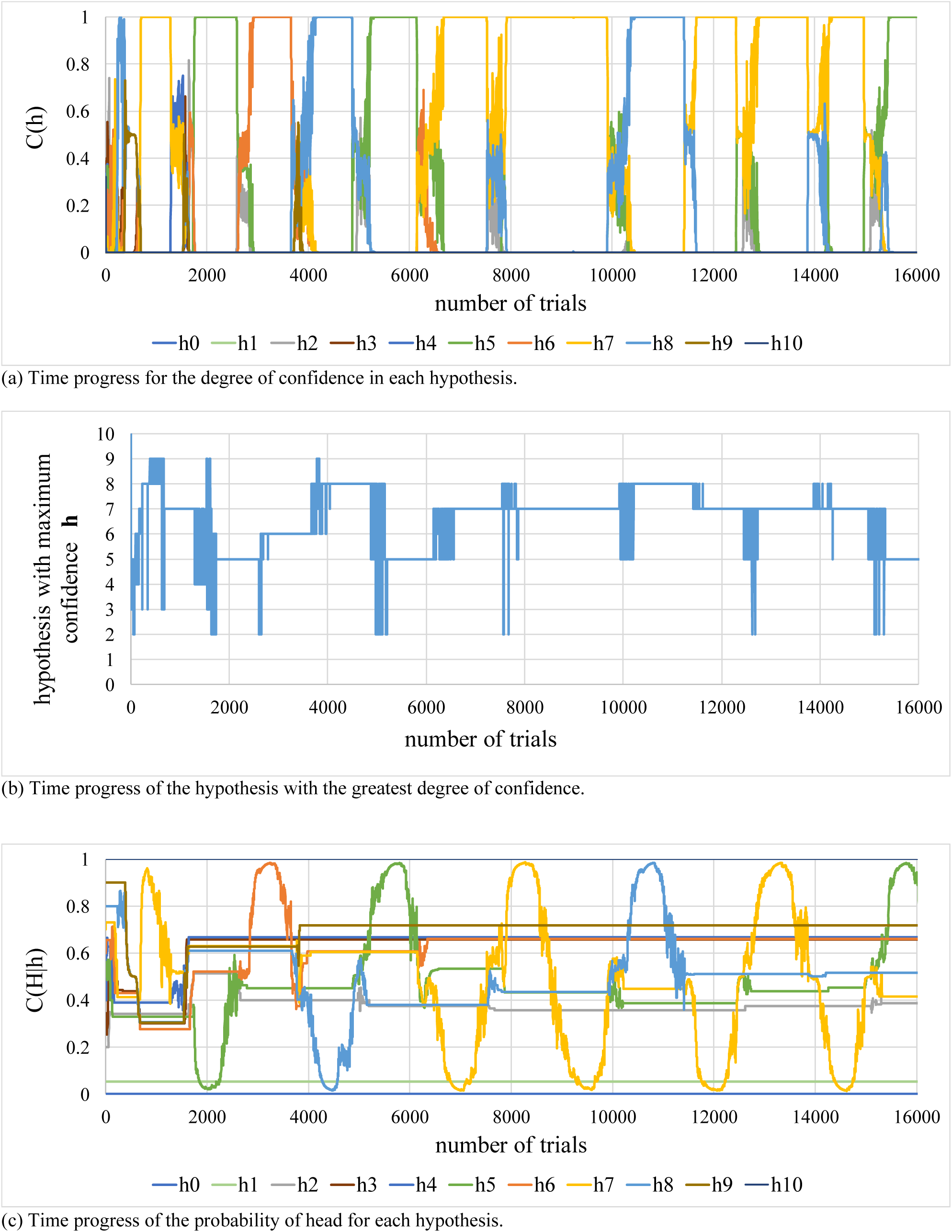
Internal state of extended Bayesian inference in Task II.

**Fig. 6.**
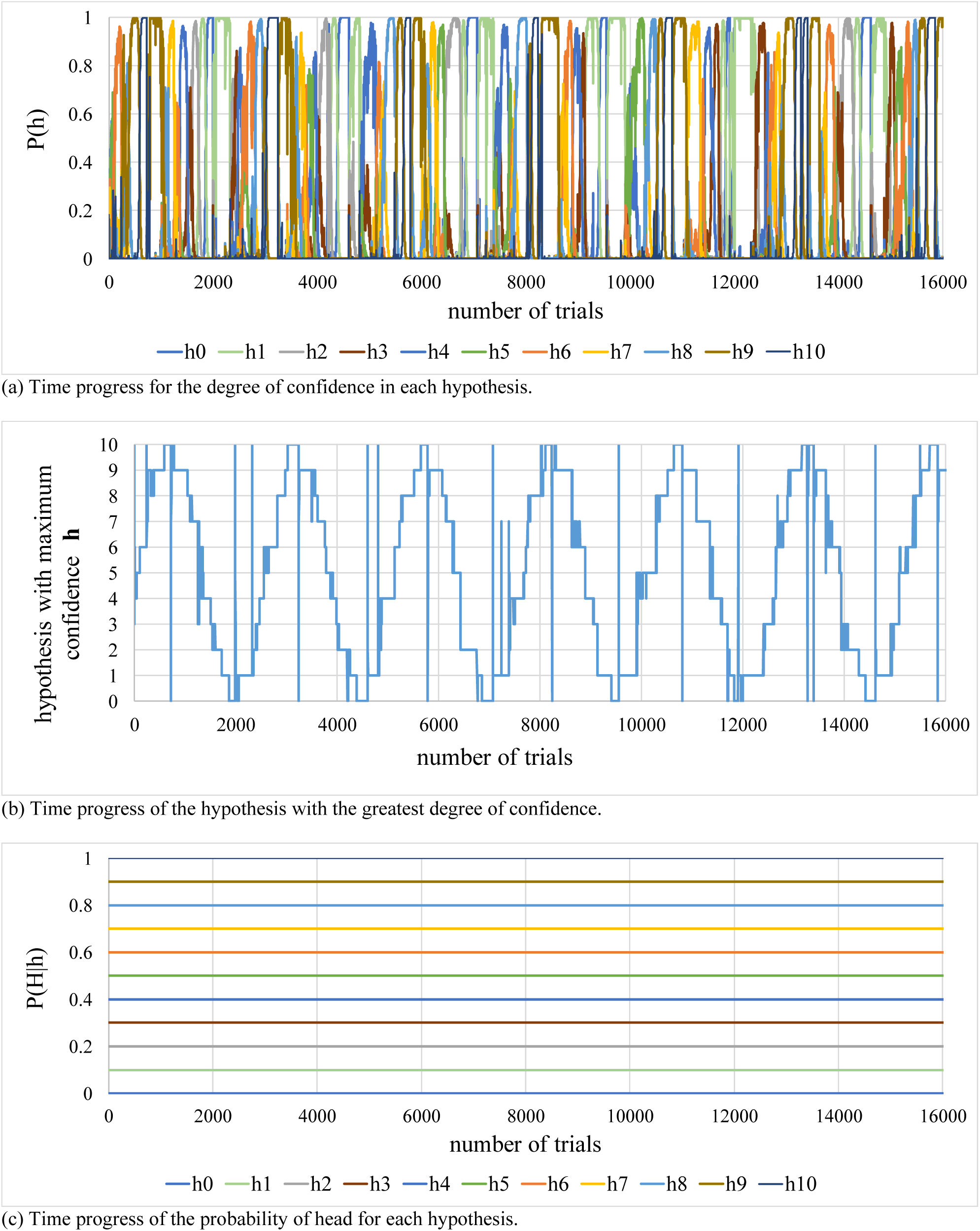
Internal state of restricted Bayesian inference in Task III.

We set *m* = −1.0. Because of this, we can rewrite formula (28) for inverse Bayesian inference as follows.

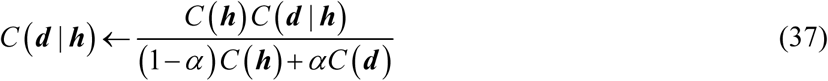

Here, we can see that the denominator on the right side is the weighted average of *C*(***h***) and *C*(***d***), if *C*(***h***) > *C*(***d***), *C*(***h***) > (1− *α*)*C*(***h***) + *αC* (***d***) so *C*(***d*** | ***h***) increases. At this point, the increment of *C*(***d*** | ***h***) is larger if the degree of confidence *C*(***h***) is higher. Conversely, if *C*(***h***) < *C*(***d***), *C*(***h***) < (1− *α*)*C*(***h***) + *αC* (***d***) so *C*(***d*** | ***h***) decreases greatly if the degree of confidence *C*(***h***) is lower. Let us turn to the analysis of the stable period where *C*(***h***) = 1. When *C*(***h***) = 1, the total sum of confidence of all hypotheses is 1, and for any hypothesis *h*_*i*_ other than ***h***, *C*(*h*_*i*_) = 0. Hence, formula (30) can be rewritten as follows.

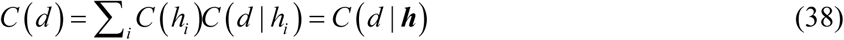

At this step, formula (37) for inverse Bayesian inference can also be rewritten as follows.

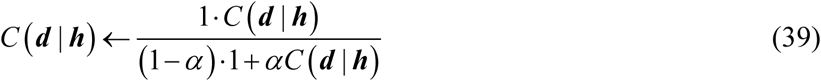

This shows the weighted harmonic average of 1 and *C*(***d*** | ***h***). Since 0 ≤ *C*(***d*** | ***h***) ≤ 1, the denominator is necessarily less than 1, and the likelihood *C*(***d*** | ***h***) increases whenever updated. In other words, when certain data are observed, the connection between the data and the hypothesis with the highest degree of confidence at that time is reinforced. Conversely, unobserved data, i.e., for *d* _*j*_ other than ***d***, *C*(*d* _*j*_ | ***h***) can be standardised using formula (29), hence decreasing by the increment amount in *C*(***d*** | ***h***). Here, the rate of increase for *C*(***d*** | ***h***) depends on *α*. When *α* = 1, regardless of the presence value of *C*(***d*** | ***h***), the right side of formula (39) is 1. As *α* gets smaller, the increase rate lowers, and when *α* = 0, it coincides with Bayesian inference and *C*(***d*** | ***h***) becomes invariable.

These considerations suggest that *α* gets larger according to increase of updates to the model. In this sense, we can say that formula (39) for inverse Bayesian inference during the steady period represents a process of learning, and *α* corresponds to the rate of learning. With respect to the portion that corresponds to Bayesian inference, suppose *m* = −1.0 in the formula (25), then we can rewrite it as:

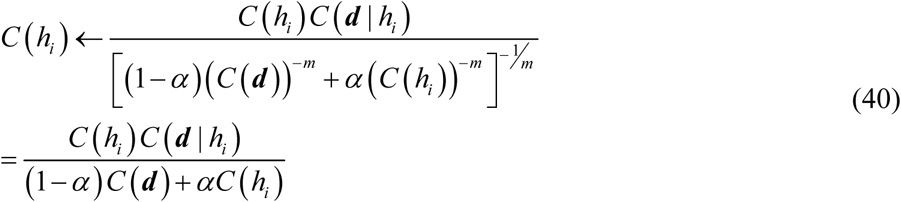

Through careful observation of this formula we can note that *α* becomes larger, while *C*(***d***), i.e., the effect of observation data, gets smaller. Where *α* = 1, the denominator disappears so *C*(*h*_*i*_) in the numerator and denominator is cancelled, and the formula can be expressed as follows.

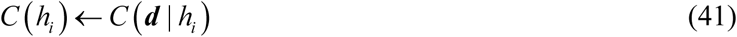

This means that when *α* = 1, the degree of confidence in each hypothesis *C*(*h*_*i*_) does not even consider the past observation history and is seen to be identical to the likelihood at that point in time. In contrast, when *α* = 0, the present formula coincides with the Bayesian inference expressed in formula (36).

A comparison of formula (36) with formula (41) reveals that their difference lies in the presence or absence of *C*(*h*_*i*_) on the right side. In this sense, the difference between them can be said to lie in the extent to which they accept history in order to determine degree of confidence.

As shown above, in extended Bayesian inference, symmetry bias plays two roles. First, the strength of symmetry bias indicates the rate of learning in portions of inverse Bayesian inference. In other words, the stronger the symmetry bias, the greater the degree of modification to the hypothesis model based on observational data. Second, the strength of the symmetry bias indicates how much the model takes into account past history in portions of Bayesian inference. In other words, as symmetry bias gets stronger, confidence in each hypothesis is updated based solely on more recent observational data.

## 4 Discussion and conclusions

In this study, we first proposed extended confidence model as a causal induction model. Then, we formulated an inference model that incorporates the said causal induction model into Bayesian inference. We noticed that this inference model necessarily involves the inverse Bayesian inference, which allows for flexibility to handle unsteady situations where the inference target changes from time to time. Finally, we demonstrated how this model can work well with unknown situations by forming new hypotheses through inverse Bayesian inference.

The causal induction model that we are proposing has two parameters that control the strength of symmetry bias. Conditional probability and causal induction models like *DFH* and *pARIs* can be shown to be special cases where particular values are assigned to parameters in our model. In other words, using the proposed model allows us to seamlessly express degree of confidence in those statements of the forms “if P then Q,” which stand for prediction, as well as those in the form of “P therefore Q,” which stand for causal relation in a single model. As shown in the appendix, the results of the meta-analysis of causal induction experiments revealed that the proper incorporation of symmetry bias in the proposed model allows it to replicate human judgment with high accuracy. However, it is known that the interpretation of conditionals largely changes depending on the type and the contents of the conditionals, as well as subject’s age (Gauffroy and Barrouillet, 2009). Further studies are necessary to determine how parameters change according to type and contents of conditionals, as well as developmental stage.

Parameter *α* that denote the strength of symmetry bias indicate, in the case of extended Bayesian inference, the strength of inverse Bayesian inference, that is, the rate of learning. At the same time, it indicates the degree to which the history is taken into account when updating the degree of confidence in each hypothesis, i.e., discounting rate. In this way, learning and inference become interlocked via the parameter that denote the strength of symmetry bias in such way that it takes account of only more recent observational data in inference as the rate of learning becomes greater.

Suppose that we have knowledge of the kinds of emotions another person has and how they are expressed. Of course, one cannot attain complete knowledge because individuals’ mental states are considered to be private to some degree. When estimating based on incomplete knowledge that a person is pleased, if his/her facial expression suddenly changes, one may think that his/her emotion has changed or that this is another way of expressing pleasure. The former is an inference based on knowledge and corresponds to Bayesian inference. The latter, on the other hand, is a modification of knowledge or an addition of new knowledge and it corresponds to inverse Bayesian inference.

Bayesian inference is effective in steady circumstances like Task I where the probability of head landing is fixed. Both Bayesian inference with *ε* > 0.0 and extended Bayesian inference can handle unsteady circumstances. However, they differ in the way they handle them: Bayesian inference handles them by replacing the hypothesis with another hypothesis while the model remains fixed. Hence, it cannot handle unknown situations where the inference target is not included in the repertoire of the model as in Task III. In contrast, since extended Bayesian inference includes the functionality of inverse Bayesian inference, even when a model for the probability of landing heads greater than 0.5 has not been given in advance, such as in Task III, it is able to handle the unknown situation well by modifying the model based on observational data.

As a framework similar to extended Bayesian inference, there are methods such as expectation-maximization (EM) algorithm and K-means. The EM algorithm is a method for obtaining the maximum likelihood estimate in a hidden variable model (Bishop, 2006) and often used for the mixture models or latent topic models such as latent Dirichlet allocation (Blei et al., 2003). K-means is the non-stochastic version of the EM algorithm (Bishop, 2006). However, in the EM algorithm, it is necessary to provide all observation data at one time. In practice, there are cases where processing must be performed sequentially each time data is observed. Various online algorithms have been proposed to deal with this situation (Neal and Hinton, 1998; Sato et al., 2000; Yamanishi et al., 2004). Comparison with these algorithms is a topic for the future.

In the simulation mentioned in this paper, parameter *α* that represents the strength of symmetry was given externally. Smaller *α* works well under steady circumstances, while larger *α* works well under highly variable circumstances. In the future, it would be interesting to contrive a system that can autonomously control these parameters. A limitation is that the proposed extended Bayesian inference can only handle discrete probability distributions. In the future, we will expand the model to handle continuous distributions such as normal distribution and apply to anomaly detection and change point detection, for example, detection of a disease sign.

## Abbreviations

(*DFH*): dual-factor heuristics
(pARIs): proportion of assumed-to-be rare instances;
(HM): harmonic mean;
(EM): expectation-maximization

## Acknowledgements

This research is partially supported by the Center of Innovation Program from the Japan Science and Technology Agency, JST and JSPS KAKENHI Grant Numbers JP16K01408.

## Author contributions

S. S. conceived the extended confidence model and the extended Bayesian inference, conducted the meta-analysis and the simulation, wrote the manuscript. U. C. revised it. T.T collected the data for the meta-analysis. N. M., K. S., U. C., T. T., P.Y.G., Y. N. and S. M. contributed to the interpretation of the study findings. All authors participated in the editing and revision of the final version of the manuscript.

## Appendix

### Evaluation of descriptive validity of extended confidence model

We perform parameter analysis by comparing extended confidence model with human experimental data obtained from literature (Anderson and Sheu, 1995; Buehner et al., 2003; Hattori and Oaksford, 2007; Lober and Shanks, 2000; White, 2003). In order to evaluate the descriptive validity of various models including the *DFH* model, Hattori and Oaksford (2007) performed a meta-analysis using data from eight types of causal induction experiments. To test the descriptive performance of the extended confidence model, we also performed the meta-analysis using the same datasets as Hattori and Oaksford (2007).

Generally, in a simple causal induction experiment, participants are given four types of co-occurrence information concerning the cause *c* and the effect *e* (Table 1). Then, they are asked to assess subjectively the strength of the causal relation between *c* and *e* using a number from 0 to 100. To measure each model’s fit to the data, we calculated the determination coefficient *R*^2^ from the pair of participants’ mean ratings of causal strength and the estimated value of each model computed from the same co-occurrence information given to the participants in the experiments. The experiment data used in the meta-analysis consists of the experiment I from (Anderson and Sheu, 1995), experiments I and III from (Buehner et al., 2003), experiments I and II from (Hattori and Oaksford, 2007), experiments I and III from (Lober and Shanks, 2000), and experiment II and VI from (White, 2003). The experiment data above will be abbreviated as AS95, BCC03.1, BCC03.3, HO07.1, HO07.2, LS00, W03.2, and W03.6 respectively. The experiments involving human participants described in this paper were not carried out by us, but we only calculated the coefficient of determination using the numerical data described in the cited papers. Therefore, we did not seek approval from the ethics committee by ourselves.

In AS95, forty graduate and undergraduate students were recruited. They were given co-occurrence information about the presence or absence of drug treatment and the presence or absence of the side effects. The subjects were asked to judge a number of problems, and each problem involved a sequence of instances of these four information types. The frequencies of each information type varied from problem to problem. At the end of a problem, the subjects were asked to enter a number from 0 to 100 that best reflected their judgment of the drugs causing the side effects.

In BCC03.1, 109 undergraduate students were recruited and divided into two groups (preventive group and generative group). In preventive group, they evaluated how effectively each vaccine prevented the corresponding disease by giving a rating on a scale from 0 (the vaccine does not prevent the disease at all) to 100 (the vaccine prevents the disease every time). They also evaluated the influence of ray exposure on the mutation of viruses in generative group.

Thirty-one undergraduate students participated in BCC03.3. With regard to the side effects of drugs that reduce allergy, participants determined whether there were side effects of headache and, if so, assessed the causal strength between the drug and the headache.

In HO07.1, participants were 39 undergraduate students. They were asked to assess the strength of the causal relation between a particular type of fertiliser and plants blooming. They only observed a sequence of scenes in which fertiliser and plant blooming were either present or absent. After observing a series of situations, participants rated the subjective strength of the causal relation with a value between 0 (completely unrelated) and 100 (completely related).

In HO07.2, participants were 50 undergraduate students. In this experiment the cause was ‘drinking milk’ and the effect was ‘stomach-ache’. They judged the causal strength between drinking milk and stomach-ache according to given co-occurrence information.

In LS00, the participants of experiments 1, 2 and 3 were 27, 16, and 24 students, respectively. They assessed the extent to which a certain chemical causes a mutation in animals’ DNA using a number from between 0 and 100, where 0 indicates that the chemical does not cause mutations at all and 100 indicates that the chemical causes a mutation.

In W03.2, the participants were 40 undergraduate students. They were given information on the additives (manganese trioxide) contained in the foods a patient has eaten, and information on whether the patient has developed an allergic reaction. They were asked to judge the extent to which the statement ‘Manganese trioxide causes the allergic reaction in this patient’ was right for that patient and to write a number from 0 (zero) to 100, where 0 (zero) means that the statement is definitely not right, and 100 means that the statement is definitely right.

In W03.6. the participants were 43 first-year undergraduate students. Most features of method, including initial written instructions; format of stimulus presentations; and procedure, were the same as in W03.2. The studies differed in design, however.

Values in parameters *α* and *m* in formula (13) were shifted each with 0.05 increment in the interval [0.0,1.0] and [−2.0,2.0] and the determination coefficient *R*^2^ between assessment by participants and the estimated value by the proposed model was calculated for each pair of parameters. Table A1 shows the pair of parameters *α* and *m* at which *R*^2^ becomes maximum in each experiment. As seen in Table A1, the determination coefficients were greater than 0.9 for all experiments. *α* was around 0.5 (0.25-0.6) and did not reach 0.0, which stands for the normal conditional probability. This suggests that symmetry bias was deeply involved in causal induction. Moreover, *m* interestingly took a negative value in all experiments. This suggest people show strong awareness of causal relations only if both *P*(*e* | *c*) and its inverse *P*(*c* | *e*) are large.

**Table A1.** Performance evaluation of the extended confidence model based on the meta-analytic data from Hattori and Oaksford (2007).

|  | AS95 | BCC03.1 | BCC03.3 | HO07.1 | HO07.2 | LS00 | W03.2 | W03.6 |
| --- | --- | --- | --- | --- | --- | --- | --- | --- |
| $R^2$ | 0.97 | 0.98 | 0.99 | 0.99 | 0.99 | 0.91 | 0.97 | 1.00 |
| $\alpha$ | 0.3 | 0.55 | 0.35 | 0.6 | 0.3 | 0.55 | 0.35 | 0.25 |
| $m$ | -0.65 | -0.45 | -0.25 | -2.0 | -2.0 | -1.4 | -1.6 | -2.0 |

$R^2$ is a determination coefficient, $\alpha$ and $m$ are the model parameters that best fit the data (i.e. highest $R^2$ ).

**Fig. A1.**
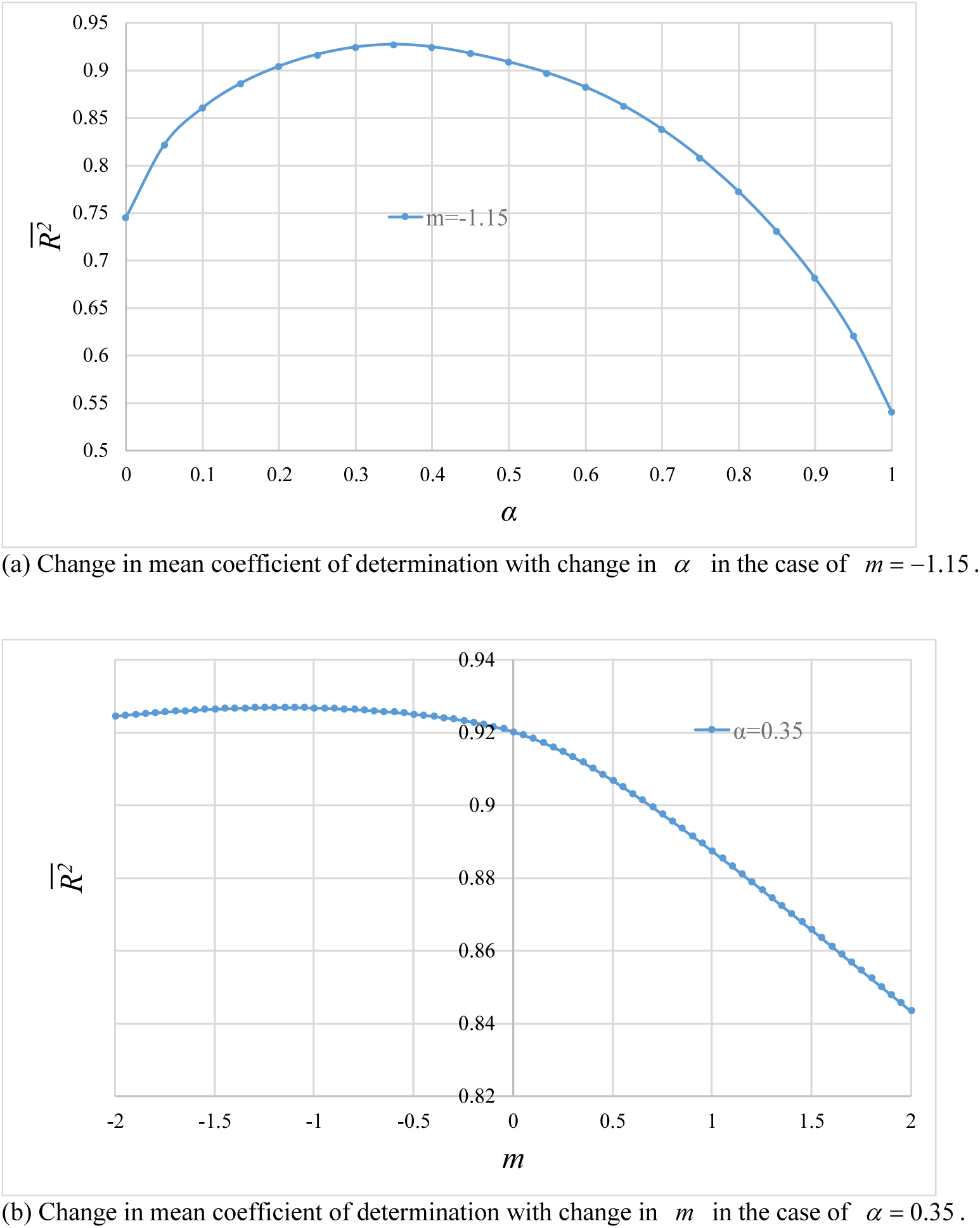
Mean coefficient of determination for each pair of parameter values. The mean value for the determination coefficient peaked when (*α,m*) = (0.35, −1.15) and the value was 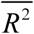 = 0.93.

In these analyses, the optimal parameter value was calculated for each experiment. In what follows, all experiments will be analysed comprehensively using common parameters. The determination coefficient when parameters have fixed values will be calculated for each experiment along with their mean. The mean value to be calculated is the weighted average using Fisher’s Z-transformation. This procedure was repeated by changing the parameter values with 0.05 increment. The mean value for the determination coefficient peaked when (*α, m*) = (0.35, −1.15) and the value was 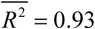. Figure A1(a) shows the change in mean coefficient of determination 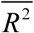 with change in *α* in the case that *m* is fixed to −1.15. The result that 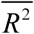 takes the maximum value when *α* = 0.35 means that human do not evaluate *P*(*e* | *c*) and *P*(*c* | *e*) equally, but rather attach importance to *P*(*e* | *c*).

Figure A1(b) shows the change in the mean coefficient of determination 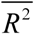 with change in *m* in the case that *α* is fixed to 0.35. In a region where *m* is positive, the value of 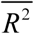 decreases rapidly as *m* increases. On the other hand, in the range where *m* is 0 or less, the value of the determination coefficient does not change much. In other words, from the viewpoint of the coefficient of determination, it can be said that it is important that *m* is 0 or less, rather than whether human’s causal induction can be expressed as the harmonic mean (*pARIs*), i.e., *m* = −1.0 or as the geometric mean (*DFH*), i.e., *m* = 0.0 of *P*(*e* | *c*) and *P*(*c* | *e*)

